# Leaf economics guides slow-fast adaptation across the geographic range of *A. thaliana*

**DOI:** 10.1101/487066

**Authors:** Kevin Sartori, François Vasseur, Cyrille Violle, Etienne Baron, Marianne Gerard, Nick Rowe, Oscar Ayala-Garay, Ananda Christophe, Laura Garcia De JalÓN, Diane Masclef, Erwan Harscouet, Maria Del Rey Granado, Agathe Chassagneux, Elena Kazakou, Denis Vile

## Abstract

The slow-fast continuum describes how resource allocation constrains life-history strategies in many organisms. In plants, it is reflected by a trade-off at the leaf level between the rate of carbon assimilation and lifespan, the so-called Leaf Economics Spectrum (LES). However, it is still unclear how the LES is connected to the slow-fast syndrome, and reflects adaptation to climate. Here, we measured growth, morpho-physiological and life-history traits at both leaf and whole-plant levels in 384 natural accessions of *Arabidopsis thaliana*. We examined the extent to which the LES continuum parallels the slow-fast continuum, and compared trait variation to neutral genetic differentiation between lineages. We found that the LES is tightly linked to variation in whole-plant functioning, relative growth rate and life history. A genetic analysis further suggested that phenotypic differentiation is linked to the evolution of different slow-fast strategies in contrasted climates. Together, our findings shed light on the physiological bases of the slow-fast continuum, and its role for plant adaptation to climate.

## Introduction

Plant populations diversify geographically as a result of adaptive (e.g., natural selection) and non-adaptive (e.g., genetic drift and isolation) processes. To understand plant evolutionary responses to current and future climate variations, it is crucial to investigate the genetic and phenotypic differentiation of plant lineages along environmental gradients. As plants cannot simultaneously optimize competing eco-physiological functions, an important question is how plant adaptation occurs under the influence of major trade-offs between traits.

The slow-fast continuum is a pervasive trade-off between resource allocation to growth, reproduction and survival, spread across the tree of life ^1^. The slow end of this continuum is characterized by slow growing, long-lived species and low reproductive output, while species at the fast end reach reproductive maturity faster and produce more offspring. In plants, the leaf economics spectrum (LES hereafter) ^2–4^, is thought to reflect the physiological basis of the slow-fast continuum ^4^. The LES arrays plant species along a continuum of leaf trait syndromes going from short-lived leaves with fast metabolism and rapid return on investment to the reverse syndrome ^3^. Core LES traits include leaf dry mass per area (LMA), leaf lifespan and net photosynthetic rate per mass unit (*A*_mass_) ^3,5–7^. Some of these leaf traits are broadly used in comparative ecology to infer whole-plant ecological strategies ^4,8–12^. However, the extent to which leaf-level resource economics reflects whole-plant physiology, performance, and ultimately fitness, is still under debate ^13^. Many processes can lead to a mismatch between LES and whole-plant functioning ^14^, including the impact of self-shading among leaves and resource allocation patterns, such as carbon investment in non-photosynthetic tissues ^15,16^. To gain insights into the robustness of the slow-fast continuum at different organizational levels, we need to examine how LES traits scale up to plant level resource-use strategies, life history and performance. However, it remains difficult to compare individual performance across species with different growth forms, phenology and dispersal strategies, which impedes a clear linkage between physiological and adaptive trade-offs ^17–19^.

The LES has been associated with differences in the ability of plants to adapt to more or less harsh environmental conditions ^4,12,20,21^: species displaying high photosynthetic, respiration and growth rates, short-lived, thin and nitrogen-rich leaves are preferentially found in nutrient-rich and/or growth-suitable climatic conditions. Those species are qualified as acquisitive in contrast to conservative species that exhibit the opposite set of traits. Despite those observations, functional ecology has no tools to test for adaptation, and empirical evidences of the adaptive value of being at one end or the other of the continuum in a given environment remain scarce (see ^22^ for review). Furthermore, sampling procedure in field observation studies often impedes to disentangle the effects of plasticity vs. genetic differentiation on the emergence of the LES ^23^. Thus, comparative studies looking for plant adaptation are at best incomplete ^22,24^, and the role of selection in shaping the LES and driving adaptation to diverse environments is hardly understood. To fill this gap, intraspecific studies are encouraged since they can take benefit from tools developed in population ecology and genetics ^22,25,26^. The LES has started to be analysed at the intraspecific level, with contrasting findings depending on the studied organism and type of study ^23,27–32^. LES relationships appeared consistent with cross-species ones when using species with broad environmental niche spectra ^31,33^ and/or broad phenotypic variability ^23^, but inconsistent when using species with narrow phenotypic (and genetic) diversity ^34^. Genetic differentiation of LES strategies has been demonstrated between populations of *Helianthus anomalus* along a 400 km rainfall gradient ^35^. However, the question whether LES diversifies because of adaptation to climate between lineages spanning large geographic distribution remains open. Overall, we still miss a comprehensive understanding of within-species LES variation and the subsequent insights they can provide to well-described interspecific patterns from an evolutionary perspective.

Adaptation requires the fixation of beneficial alleles in a population, which subsequently leads to phenotypic variation and differential response to the environment. However, neutral alleles can also vary in abundance due to the effect of genetic drift. By comparing genetic and phenotypic differentiation between populations or lineages, Qst-Fst provides a powerful tool to infer adaptation in polygenic quantitative traits such as LES traits^36^. It notably enables to decipher neutral and adaptive demographic processes at the origin of phenotypic diversity within species. Indeed, Qst-Fst comparisons are based on the computation of genetic (Fst) and phenotypic (Qst) differentiation between populations or lineages. Hence, Qst values above neutral Fst are interpreted as a signature of diversifying selection on the underlying trait. For instance, Qst-Fst comparisons have been successfully used in the species *Campanula rotundifolia, Arrhenatherum elatius* and *Quercus oleoides* to investigate the role of selection in the diversification of life-history traits, growth strategies or drought resistance between lineages at both local and global scales ^37–39^. Thus, Qst-Fst comparison appears promising in the perspective of examining how phenotypic trade-offs trigger local adaptation along geographical and environmental gradients^35^. This method is expected to be particularly powerful in model species, with the help of modern genomics such as high-throughput genotyping and sequencing ^40,36^.

The species *Arabidopsis thaliana* has been widely used in molecular biology, cell biology and quantitative genetics. Thanks to the efforts to characterize the genetic diversity in this species^41–44^, it is also a model in population dynamics ^45^ and evolutionary ecology ^46^. For instance, the genetic determinism of *A. thaliana* life history has been extensively studied, notably with the discovery of genes that control major developmental transitions such as flowering time ^47^. Allelic variation in these genes appears to be adaptive to climatic and altitudinal gradients ^48^. A recent study in *A. thaliana* supports the hypothetic link between life history variation and the LES, highlighted by strong genetic correlations between these traits ^28,49^. However this analysis was performed on recombinants inbred lines used for genetic mapping. Made of artificial crosses, they preclude examining the relationships between LES and the natural environment. Interestingly, *A. thaliana* has recently gained a renewed interest in functional ecology and biogeography^50,51^, notably due to the large panel of natural accessions that have been collected from contrasting climates, and genotyped at high density (e.g. ^41–44^). As genetic data in *A. thaliana* allows an unprecedented large-scale analysis of genetic variation between populations and lineages, this species is promising to investigate the extent of intraspecific diversity in LES traits and its role for adaptation to contrasted climates.

In this study, we explored the evolutionary bases of LES variation using a pan-European collection of 384 natural accessions from the RegMap panel ^43^. Specifically, we investigated whether plant adaptation to various climates is associated with genetic differentiation along the slow-fast continuum. To test this hypothesis, we first examined how the LES shapes phenotypic diversity across contrasted accessions of *A. thaliana*, and tested whether LES traits scale up to plant level resource-use strategies, life history and performance. Secondly, we took benefit from the large genomic information available in *A. thaliana* to evaluate with Qst-Fst comparisons to what extend LES differences between lineages are attributable to adaptive processes such as adaptation to contrasted climates.

## Results

### Geographic clustering of *A. thaliana* lineages

The range of biomes experienced by the sampled genotypes covers temperate grasslands, deserts, woodlands-shrublands and temperate forests (Fig. 1b). Using the 250K SNPs data available from Horton et al. ^43^, we performed a genetic clustering of the genotype set. A cross-validation for different numbers of clusters (k = 3 to k = 11) showed that our set of genotypes can be separated into six groups representative of different genetic lineages (cross validation error = 0.89). These lineages were moderately differentiated (mean F_ST_ = 0.11), geographically (Fig. 1a) as well as in the Whittaker’s biome classification (Fig. 1b). The analysis revealed the existence of two genetic groups exclusively located in France in our sample (French 1 and French 2 hereafter) of 73 and 45 genotypes, respectively. Among the 15 genotypes of the third group, seven were defined as North Swedish in the 1001 genomes dataset ^44^. Consistently, the 15 “Swedish” genotypes, although not all in Sweden (Fig. 1a), were mainly located in cold environments and woodland-shrubland in Whittaker’s classification (Fig. 1b). We considered genotypes from group 4 as “Central European” (Fig. 1a), typically living at intermediate temperatures and rainfall (Fig. 1b). 68 genotypes composed the group 5, all located in Western Europe (Fig. 1a), in a range of relatively warm environments with intermediate rainfall. Finally, only five genotypes composed group 6, all collected in USA (“American” genotypes hereafter).

**Figure 1.**
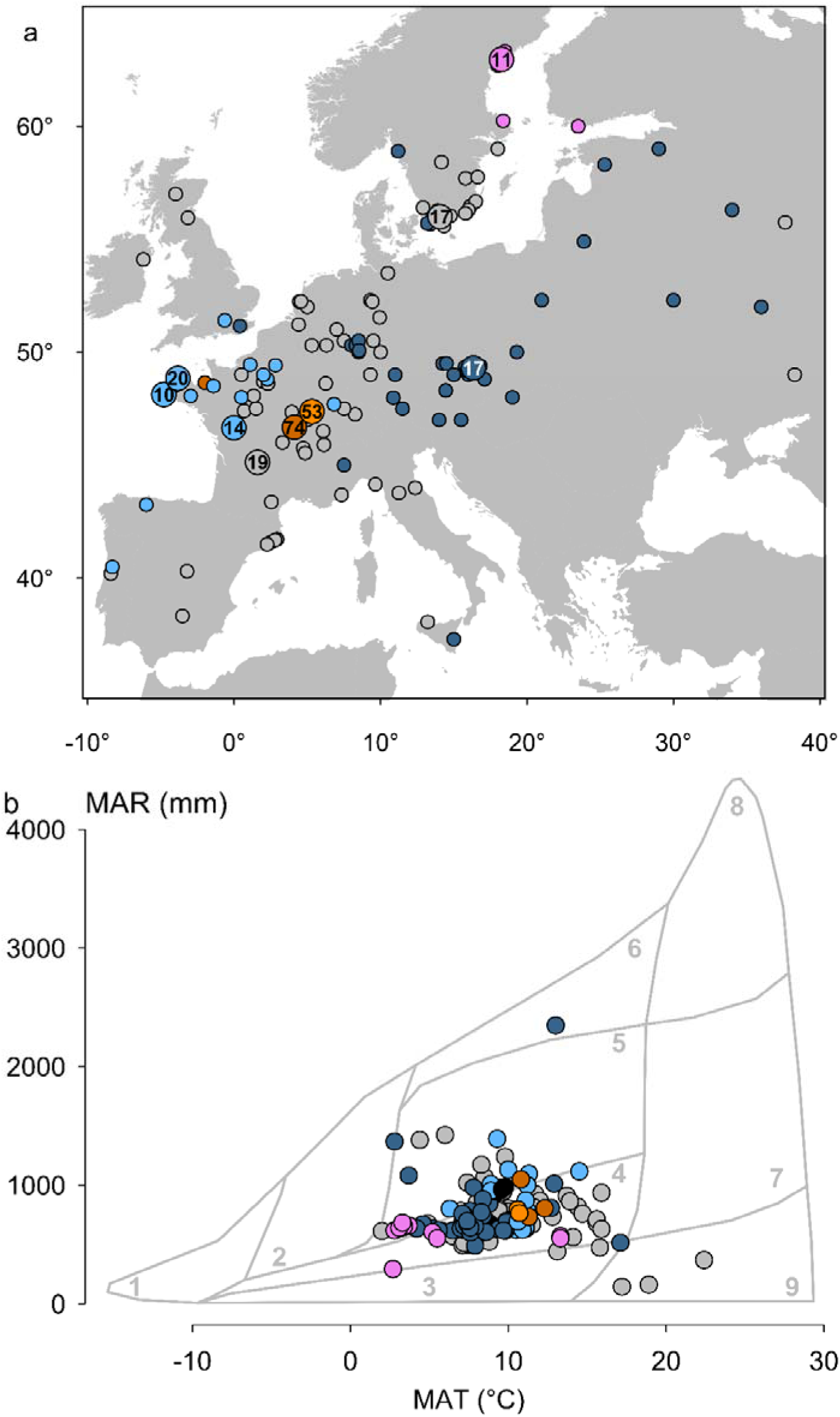
Location and climatic conditions of the genotype collecting sites. **(a)** Distribution of the 384 natural genotypes used in this study. The small points represent the collecting sites of genotypes and bigger points give the number of collecting sites overlapped at these positions. The colours represent the six genetic groups: Admixed (grey), French1 (brown), French2 (orange), Swedish (purple), Central Europe (dark blue), Western Europe (light blue), American (black). **(b)** Mean annual rainfall (MAR) and mean annual temperature (MAT) for the sites where genotypes were collected, in relation to major biome types of the world following Whittaker’s classification. 1-9: Tundra, Boreal forest, Temperate Grassland Desert, Woodland Shrubland, Temperate Forest, Temperate Rain Forest, Tropical Forest Savana, Tropical Rain Forest, and Desert.

### Leaf economics of *A. thaliana*

Assimilation rate was the most variable trait among the leaf economics traits in our dataset (18-fold; from 34.8 to 608.9 µmol g^-1^ s^-1^) while LMA and LLS varied 5 and 3.5 times (from 18.7 to 101 g m^-2^, and from 15 to 53.5 d), respectively. Representing the dataset within this 3-dimension space revealed that the LES constrains the covariation of *A. thaliana* leaf traits. Consistent with interspecific comparisons, genotypes are ranked from low A_mass_ and high LMA and LLS, toward high A_mass_ and low LMA and LLS (Fig. 2). A principal component analysis (PCA) showed that 78% of the covariation between these three traits was explained by a single Principal Component (Fig. S1a). Hereafter, we assigned a position along the LES for each genotype with its score on PC1. *A*_mass_ was highly negatively correlated with PC1 (r = -0.91) while LMA and LLS were positively correlated with PC1 (r = 0.93 and 0.81, respectively). Thus, high and low PC1 values are representative of genotypes located at the conservative and acquisitive side of the LES, respectively.

**Figure 2.**
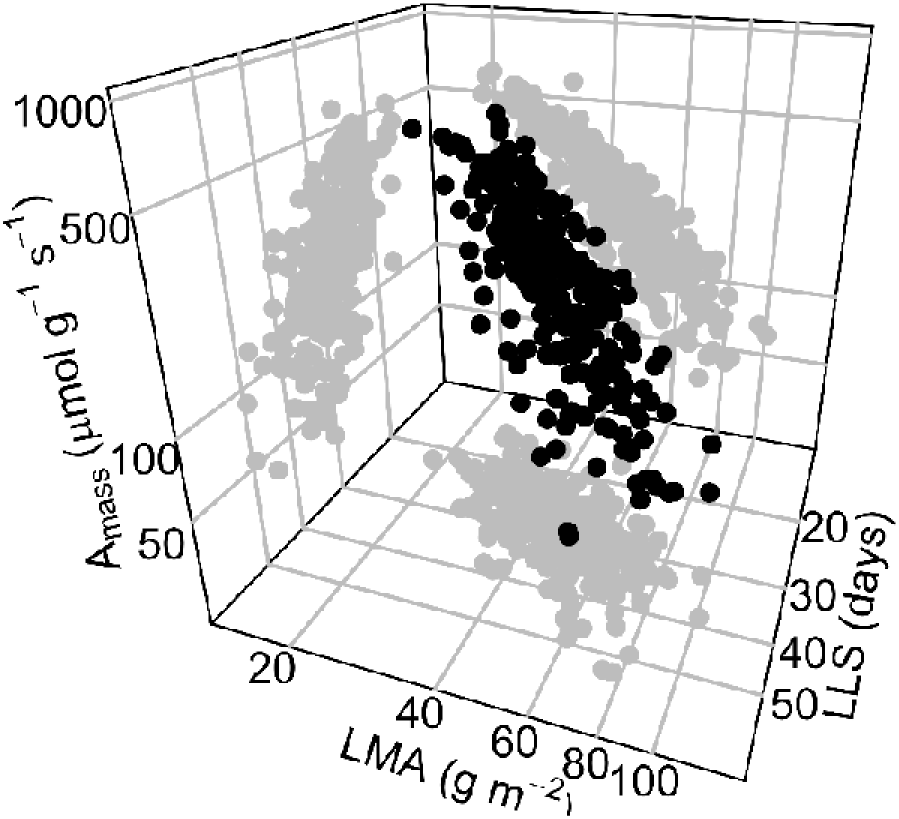
The leaf economics spectrum in *A. thaliana.* Three-way relationships among the main leaf economics traits: A_mass_, mass based assimilation rate (nmol CO_2_ g^−1^ s–^−1^); LMA, leaf mass per area (g m^-2^); LLS, leaf lifespan (d).

### From the leaf economics spectrum to the plant slow-fast continuum

Trait measurement at the plant level revealed that assimilation rate was again the most variable trait with a 68-fold variation (from 8.4 to 578.1 µmol g^-1^ s^-1^), while plant mass per area and age of maturity varied 5 times (from 17.7 to 85.4 g m^-2^ and 22 to 111 d, respectively). Standardized major axis regressions between traits measured at the leaf and plant levels were highly significant. Leaf and plant-level LMA were highly correlated (r = 0.78; P < 0.001; Fig. S1c) and the slope was close to, but significantly different from 1 (95% CI slope = [1.07, 1.17]), as well as for leaf-level and plant-level net photosynthetic rate (r = 0.84; P < 0.001; Fig. S1d, 95% CI slope = [0.79, 0.9]). Similarly, leaf life-span and plant age of maturity were significantly correlated with a slope below 1 (slope = 0.58 [0.53; 0.63], r = 0.47, P < 0.001; Fig. S1e). A single principal component explained 87% of the trait covariation at the plant level (Fig S1b) and was highly correlated with the PC1 of the PCA on leaf traits (r = 0.87, p < 0.001). Indeed, life history and performance at the plant level were strongly related to PC1, as illustrated by the correlation between relative growth rate (RGR) and PC1 (r = -0.33) and age of maturity (AM) and PC1 (r = 0.76) (Fig. 3a,b), highly significant when including or not the kinship matrix as a covariate (p < 0.001). Conservative leaf strategies are thus associated with slow plant growth and late reproduction, whereas acquisitive leaf strategies are associated with fast plant growth and early reproduction.

**Figure 3.**
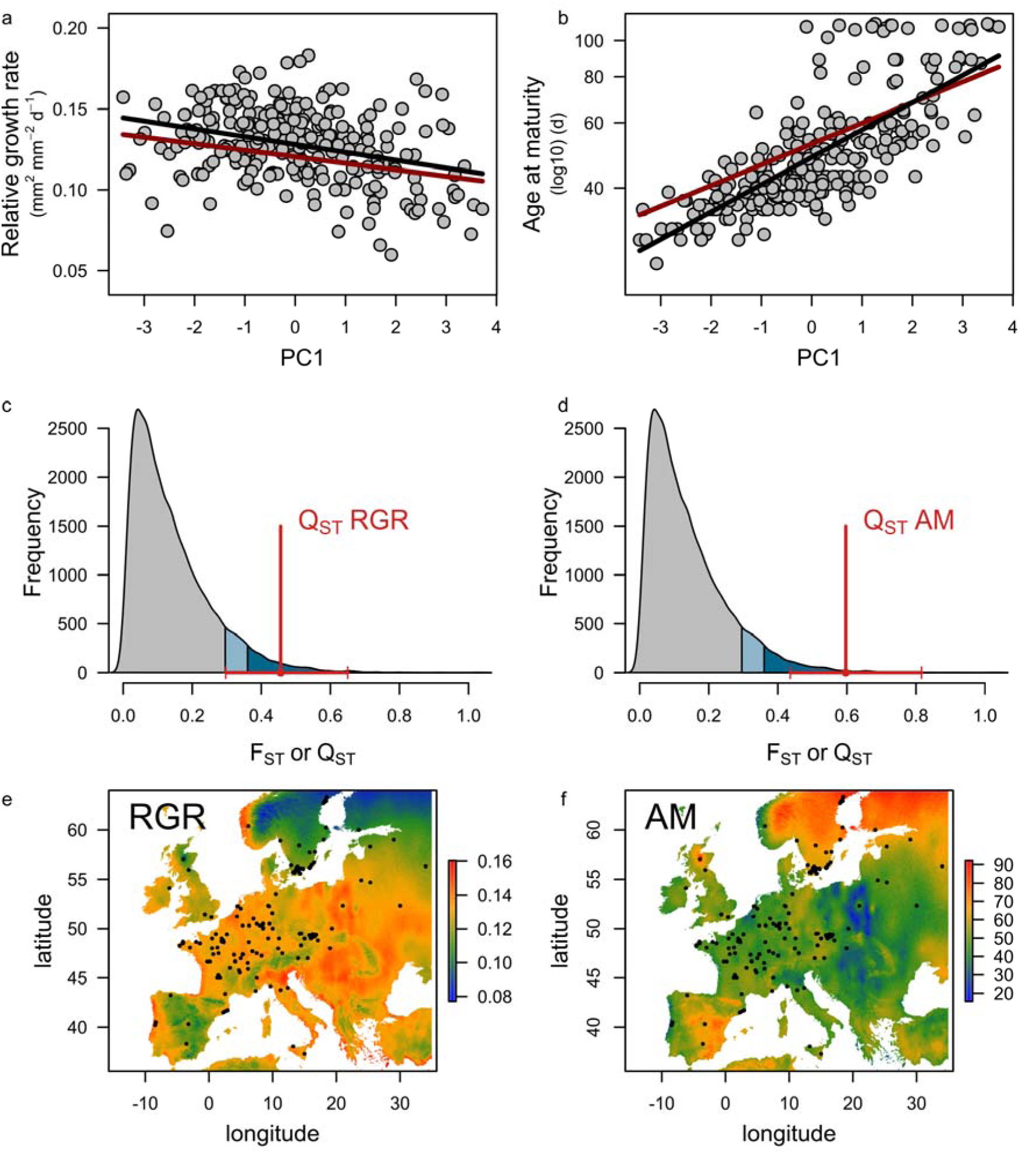
The slow-fast continuum makes the bridge between leaf economics spectrum and plant response to climate. (a) Relative growth rate (RGR, mm² mm^-2^ d^-1^) and (b) age of maturity (AM, d) as a function of the first principal component of LES traits. Regressions and phylogenetic regressions are represented by black and red lines, respectively, when significant. Q_ST_ values and 95% CI for (c) RGR and (d) AM relatively to the 95^th^ quantile (blue area) and 90^th^ quantile (light blue area) of the F_ST_ distribution (grey area). Representation of the climatic predictions for (e) RGR and (f) AM. Black dots represent genotypes collecting sites.

### Adaptation cues of the slow-fast continuum

We performed Q_ST_-F_ST_ comparisons for both RGR and AM to assess the adaptability of the slow-fast strategies. Pairwise comparisons based on the whole genetic data indicated a strong genetic divergence between French2 and the other groups (F_ST_ > 0.19), as well as between American and other groups (F_ST_ > 0.43), although the low number of accessions belonging to the American group can bias the latter. French1 group was genetically closer to Western European group (F_ST_ = 0.13) than French2 group (F_ST_ = 0.20). Interestingly, North Swedish lines showed strong phenotypic and genetic differentiation with other lineages (Fig. S2). While the average genetic differentiation between genetic groups was moderate (mean F_ST_ = 0.11), phenotypic differentiation was remarkably high for RGR and AM compared to the F_ST_ distribution. Further, the high skewness of the F_ST_ distribution (1.46) suggested that the large majority of SNPs can be considered as neutral while the RGR and AM Q_ST_ fell on the upper 5% of the F_ST_ distribution (Fig. 3c,d). We performed a parametric bootstrap analysis to generate confidence intervals around Q_ST_ values and showed that the lowest estimates of the relative growth rate (age of maturity) Q_ST_ fell on the upper 10% (5%) of the F_ST_ distribution. Similarly, we showed that all LES traits but leaf lifespan, exhibited extreme Q_ST_ values (Fig. S1f). This suggests that slow-fast traits at both leaf and plant levels behaved like outlier variants that strongly diverged between lineages due to the effect of diversifying selection.

### Climatic drivers of *A. thaliana* phenotypes

We investigated whether 19 climatic variables (www.worldclim.org/bioclim) at the collecting sites of the genotypes explain *A. thaliana* slow-fast strategies. Stepwise regressions revealed that both RGR and AM are best predicted by a subset of climatic variables (r² ~ 0.16 in both cases, p < 0.001). More precisely, the annual mean temperature, annual mean precipitation, and temperature seasonality of the collecting sites had a positive (negative) effect on RGR (AM), while the mean temperature of the warmest quarter had a negative (positive) effect. The best predictor model of RGR (AM) variation included as well the minimal temperature of the coldest month (positive effect), and the best predictor model of the AM included the maximum temperature of the warmest month (negative effect). Using the WorldClim database, we predicted a theoretical map of the slow-fast continuum for *A. thaliana*. We showed that slow strategies, characterized by slow growth and late reproduction, were favoured in North Europe and Central East of Spain and in the highest European reliefs. In addition, fast strategies characterized by fast growth and early reproduction were found in Central Europe and near the coasts.

## Discussion

The comparison of multiple species based on a few traits is the historical approach of functional ecology ^52^. While fruitful ^12^, such an approach impedes a deeper investigation of how evolutionary forces and trade-offs operate together to shape the observable phenotypic diversity ^24,26^. Notably, several trait-trait covariations have been discussed in functional ecology in the light of trade-off theories. One of the most prominent phenotypic pattern discussed in the last decades, the so-called Leaf Economics Spectrum (LES), is thought to reflect a trade-off between metabolic rate and lifespan at the leaf level ^3,5,53,54^. Plant species that exhibit long-lived leaves have been referred to nutrient conservative species. They optimize long-term carbon gain and extended nutrient time residence, as well as nutrient use efficiency ^55^. By contrast species with short-lived leaves sacrifice nutrient retention to maximize the rate of carbon fixation. The LES is expected to reflect an adaptive trade-off between fast and slow growth strategies across plant species ^4^. Two assumptions underline this assertion: (i) the negative correlation between leaf photosynthetic rate and leaf lifespan is translated into a negative correlation between plant growth rate and the duration of the life cycle, (ii) particular combinations of slow-fast traits are selected in different environments. Both assumptions are difficult to test at the interspecific level. This has generated a living debate about the evolutionary causes of the LES ^22,28,56–59^. Taking benefit from a large collection of sequenced ecotypes in a model species, our results show that LES traits are correlated with slow-fast strategies at the plant level, and that trait divergence between genetic lineages is non-neutral. This supports the idea that plant populations evolve different slow-fast strategies along with different LES traits in order to adapt to contrasting climates.

We showed that LES trait correlations in *A. thaliana* follow the interspecific pattern ^3,5^: individuals that invest a large amount of biomass per unit leaf area have a lower leaf assimilation rate and a longer leaf lifespan than plants that invest less biomass per unit leaf area. Moreover, the economics spectrum is conserved when scaling from leaf to whole-plant traits. This gives strong support to the idea that a trait value obtained on a single leaf using a standardized method, reflects the average phenotypic value expressed by all the leaves of an individual plant ^60,61^. Furthermore, our results showed that carbon economy at the leaf level is connected to the slow-fast strategies at the plant level: the LES explained variations in growth rate and age of maturity in *A. thaliana*, two core traits of the slow-fast continuum. This suggests that the strong coordination between leaf assimilation rate, leaf lifespan and LMA act as a guide for the slow-fast strategies at the whole plant level. However, the correlations between leaf-level and whole-plant traits are presumably strongly variable between species. It is notably expected to be weaker in woody species because of the varying proportion of non-photosynthesizing tissues ^62^. Our results illustrate this statement; the assimilation rate decreases faster and the mass per unit area increases faster at the plant level than the leaf level. Similarly, leaf lifespan varied less and increased more slowly than plant age of maturity, which gives room for a decoupling between leaf and plant life history. Nonetheless, the ranking between genotypes is conserved and allows a scaling of strategies. Although our study is a first step in this direction, further explorations of how leaf level trade-offs constrains plant functioning in herbaceous and woody species are needed. In this perspective, several studies reported positive covariations between leaf carbon assimilation rate and plant relative growth rate in young trees ^63,64^.

Using F_ST_-Q_ST_ comparisons, we demonstrated how a whole plant syndrome guides the *A. thaliana* phenotypic differentiation across contrasted climates. The predicted distribution of slow-fast strategies across Europe revealed differential selection between roughly Norway, Sweden and Spain on one side, and central and Western Europe on the other. Selection for slow ecotypes toward higher latitude in *A. thaliana*, specifically in North Swedish accessions, is supported by previous findings on flowering time ^65,66^. More surprisingly however, our results suggest that similar trait combinations representative of slow strategies are selected in two contrasted climates: Spain and Scandinavia, which are at the opposite edges of the *A. thaliana* latitudinal range. This clustering of *A. thaliana* genotypes echoes a recent study showing drought related allele fixation in both Scandinavian and Spanish *A. thaliana* populations ^67^. If we consider together the absence of significant effect of the kinship matrix on trait-trait relationships tested, the globally low average differentiation between genetic groups (F_ST_ = 0.11), and the phenotypic similarity observed at two distant locations, it suggests that genetic determinism of slow strategies as well as phenotypic differentiation could have occurred by convergence through adaptive processes. Thus, slow strategies could be selected in response to water limitation in regions from nonetheless very different climates: low average temperature at Scandinavian sites, high altitude at Spanish sites (between 300 and 1100 m). Interspecific studies at global scale revealed a negative relationship between conservative strategies and rainfall ^3,68^, possibly linked to a higher investment in cell wall complex macromolecules to face drought stress ^69^. Large-scale interspecific studies also reported a bias toward acquisitive strategies with increasing temperature in herbaceous species ^70,71^. Together, this suggests a general selection pressure for slow strategies in dry environments, opposed to selection for fast strategies in non-stressing environments ^23^.

Using a model species, with large collections of well-characterized genetic material, appears particularly successful to go deeper into the evolutionary underpinning of major eco-physiological trade-offs, such as the LES and the slow-fast continuum. Combined with global climatic data, our findings notably revealed the role of selection for drought on slow-fast strategies in *A. thaliana*. Next steps will be to merge approaches, and fully benefit from what a model species can provide both genetically and eco-physiologically. For instance, the climatic cues detected here despite the lack of climate data precision, is encouraging for the future of functional biogeography ^72^. There is also evidence that the connection between functional trait and environmental adaptation requires a better characterisation of plant fitness through demographic measures^26^. Comparative studies integrating demographic approach at population level are promising to understand how selection and macro-ecological gradient shape the evolutionary responses of plants to climate variation ^24,26^.

## Materials and methods

### Plant material

We used a total of 384 natural genotypes of *Arabidopsis thaliana* L. Heynh sampled from the worldwide lines of the RegMap population (http://bergelson.uchicago.edu/wp-content/uploads/2015/04/Justins-360-lines.xls), which were genotyped for 250K bi-allelic SNPs (Horton *et al.*, 2012).

### Growth conditions

Phenotype characterization was performed under controlled conditions in the high-throughput PHENOPSIS phenotyping platform (Granier *et al.* 2006) to track daily growth. Seeds were kept in the dark at 4 °C for at least one week before sowing. Four to six seeds per genotype were sown at the soil surface in 225 ml pots filled with a 1:1 (v:v) mixture of loamy soil and organic compost (Neuhaus N2). The soil surface was moistened with one-tenth strength Hoagland solution, and pots were kept in the dark during 48 h under controlled environmental conditions (20 °C, 70% air relative humidity). Then, pots were placed in the PHENOPSIS growth chamber at 20 °C, 12 h photoperiod, 70 % relative humidity, 175 µmol m^-2^ s^-1^ PPFD. Pots were sprayed with deionized water three times per day until germination, and then soil water content was adjusted to 0.35 g H_2_O g^-1^ dry soil (–0.07 MPa soil water potential) to ensure optimal growth (Aguirrezábal *et al.* 2006; Vile *et al.* 2012). After emergence of the fourth leaf, one plant individual was left in each pot.

### Measurements of plant traits

In order to standardize measurements at an ontogenic stage for all genotypes, all traits were quantified when flower buds were macroscopically visible (i.e. bolting stage), and leaf traits were measured on the last adult leaf, fully exposed to light.

Net photosynthetic rate, relative expansion rate, lifespan, vegetative dry weight, as well as leaf area were determined for the leaf and the plant canopy. Net photosynthetic rate was measured at leaf (leaf *A*, nmol CO_2_ s^−1^) and whole-plant levels (plant *A*, nmol CO_2_ s^−1^) under growing conditions using, respectively, the leaf cuvette provided with the infrared gas analyser system (CIRAS 2, PP systems, USA), and a whole-plant chamber prototype designed for *A. thaliana* by M. Dauzat (INRA, Montpellier, France) and K. J. Parkinson (PP System, UK) (see Vasseur et al. 2012). Leaf and whole-plant photosynthetic rates were both expressed on dry mass basis (leaf *A*_mass_ and plant *A*_mass_, nmol CO_2_ g^−1^ s^−1^). Due to time constraints, we measured photosynthetic rates for 319 and 348 accessions at the leaf and whole-plant levels (306 in common), respectively. We estimated the age of maturity by the day length from germination to the appearance of the flower bud. Then, plants were harvested, and individual fresh weight was determined. The leaf used for photosynthetic measurements was identified and processed separately, and detached rosettes were kept in deionised water at 4 °C for 24 h, and water-saturated weight was determined. Individual leaves were then attached to a sheet of paper and scanned for subsequent determination of the leaf number and total leaf area using ImageJ (Schneider *et al.*, 2012). Dry weight of laminas and petioles were obtained after drying for 72 h at 65 °C. Rosette dry weight was expressed as the sum of lamina and petiole dry weights. Leaf mass per area was both calculated for the leaf used for photosynthetic measurements (LMA, g m^-2^) and for the whole-rosette (plant LMA, g m^-2^) as the ratio of lamina dry mass to lamina area. Relative growth rate (RGR, mm² mm^-2^ d^-1^) and leaf lifespan (LLS, d) were estimated from automated daily pictures of the rosettes. More precisely, a sigmoid curve was fitted to rosette area as a function of time in order to extract growth parameters, where RGR was calculated as the slope at the inflection point ^73–75^. Using daily pictures, we tracked three consecutive leaves from birth (emergence) to death (full senescence). For each plant, leaf duration was calculated as the average number of days from leaf emergence to senescence.

### F_ST_ and Q_ST_ measures

In order to perform population genetic analyses, genetic groups were identified by genetic clustering of 384 genotypes, using the 250K SNPs data available from Horton et al. ^43^. Clustering was performed with ADMIXTURE ^76^ after linkage desequilibrium pruning (*r*^2^ < in a 50 kb window with a step size of 50 SNPs) with PLINK ^77^, resulting in 24,562 independent SNPs used for subsequent analyses. Following the same approach as the 1001 genomes project ^44^, we assigned each genotype to a group if more than 60% of its genome derived from the corresponding cluster. The 123 accessions not matching this criterion were labelled ‘‘Admixed’’ and were not used for the F_ST_ and Q_ST_ calculation. The groups genetically defined were also geographically distinct. We calculated Weir and Cockerham F_ST_ value for all the 24,562 SNPs, as well as mean F_ST_ genome-wide. We also calculated Q_ST_, an analogue of F_ST_ measure, used to estimate the diversification of quantitative traits among populations, as the between-group variance divided by the total variance for leaf and whole-plant traits. A value of Q_ST_ higher than neutral loci F_ST_ or the 95^th^ quantile of the F_ST_ distribution means that the phenotypic differentiation between populations is larger than expected by demographic events alone, in particular genetic drift, and is thus indicative of diversifying selection on traits ^36,78^. We used parametric bootstrap method to generate 95% confidence intervals (CI) around Q_ST_ values with the package MCMCglmm in R (10,000 iterations).

### Statistical analysis

Climate variables at the sampling sites of each genotype were extracted from the Worldclim database (http://www.worldclim.org/bioclim), with a 2.5 arc-minutes resolution. The effect of climatic variables on traits was tested using linear model regressions. All analyses were performed in R 3.4.1 2017 (R Core Team, 2017). Whittaker biomes were plotted using the BIOMEplot function provided by G. Kunstler (https://rdrr.io/github/kunstler/BIOMEplot/src/R/biomes-plot.R). Principal component analysis (PCA) was performed using the package factominer. The package nlme was used to perform linear models and phylogenetic generalized least squares regressions. SMA regressions were performed using the package smatr ^79^, and phylogenetic SMA regressions using the Phyl.RMA function of the Phytools package. We performed phylogenetic regressions (LM and SMA) including a relatedness matrix as covariance matrix, obtained after running the PLINK--make-rel command across the 214,051 SNPs from the RegMap data.

## Supporting information

## Acknowledgments

We thank Myriam Dauzat for her technical assistance during trait measurement and environmental control of the PHENOPSIS. This work was supported by INRA CNRS and the European Research Council (ERC) (‘CONSTRAINTS’: grant ERC-StG-2014-639706-CONSTRAINTS to CV). This publication has been written with the support of the Agreenskills fellowship programme which has received funding from the EU’s Seventh Framework Programme under the agreement N° FP7-609398 (Agreenskills contract 3215 ‘AraBreed’ to FV).

## Author contribution

DV, FV and CV designed the study, KS, EB, MG, OA-G, AC, LGDJ, DM, EH, MDRG and AC conducted the experiments. KS and FV performed statistical analyses. KS wrote the first draft of the manuscript, and all authors contributed to revisions.

## Data availability statement

The authors agree that the data supporting the results will be archived in an appropriate public repository and the links and identifiers will be included within the article.

