## Supplementary material for "Leaf economics guides slow-fast adaptation across the geographic range of *A. thaliana*"

### Supplemental Information

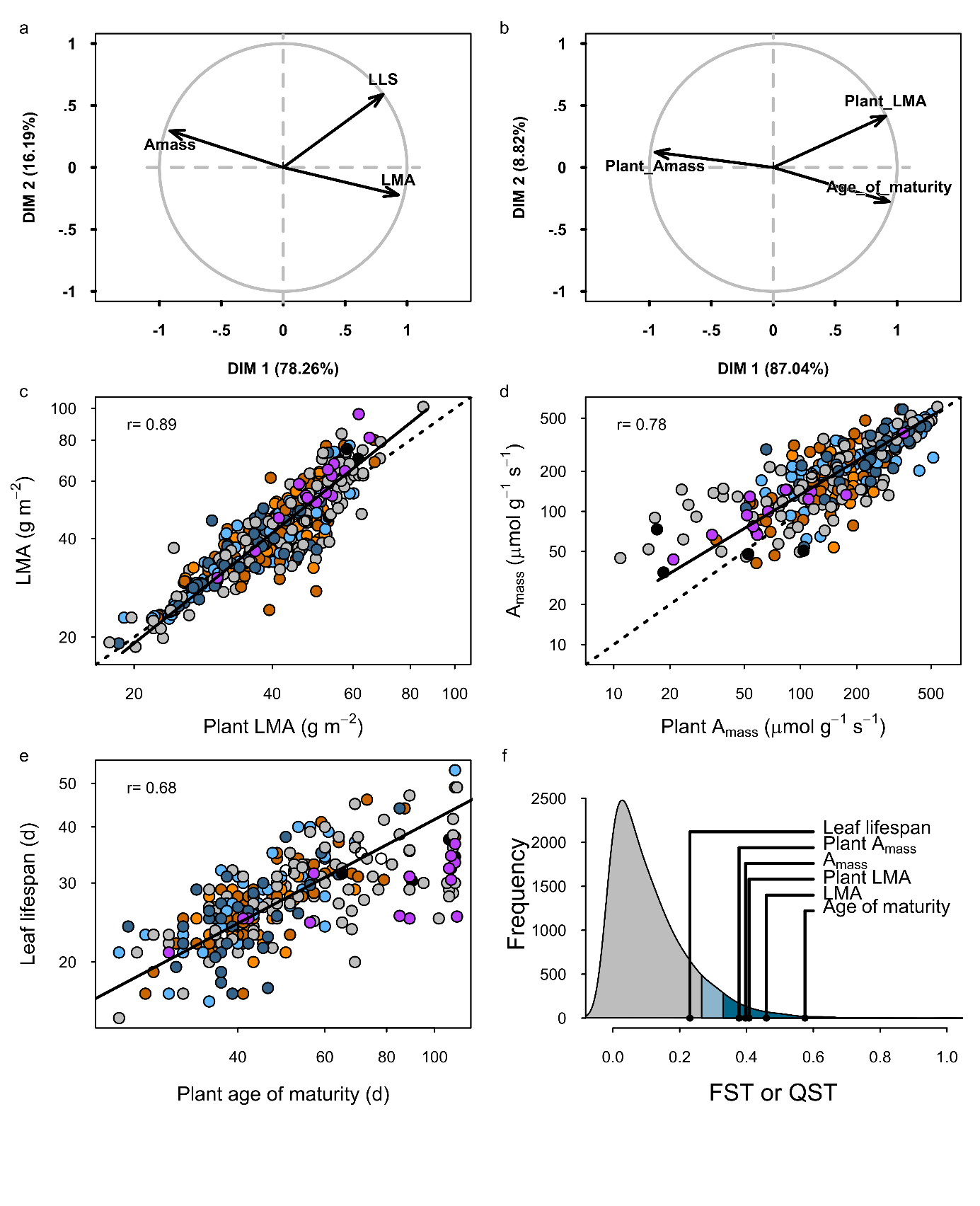

**Figure S1: Leaf economics traits scale from leaf to plant level in *A. thaliana,* and most of them are under selection.** Principal component analysis of LES traits (Amass, Assimilation rate; LLS, leaf lifespan; LMA, leaf mass per area) at the leaf (a) and plant levels (b). Covariation between leaf and plant traits: Leaf mass per area (c), Assimilation rate (d), life history (e). Dashed lines represent the identity relation and continuous lines represent standard major axis regression when significant. Q_ST_ values and 95% CI leaf and plant economics traits relatively to the 95^th^ quantile (blue area) and 90^th^ quantile (light blue area) of the F_ST_ distribution (grey area) (f).

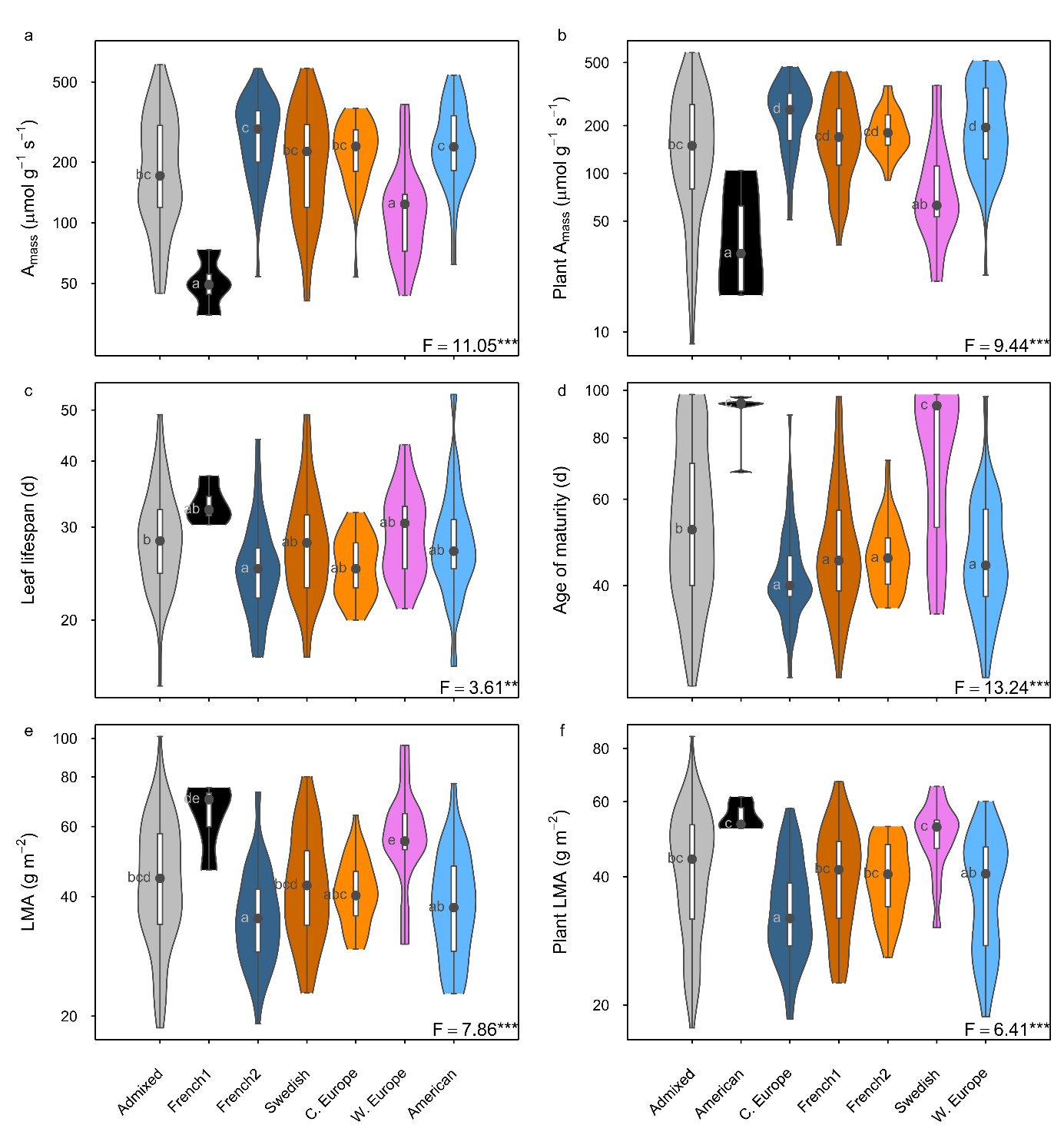

**Figure S2: Mean trait comparison between genetic groups.** Leaf (a) and plant (b) assimilation rate per unit mass (Amass), leaf (c) and plant (d) life history trait, leaf mass per area (LMA) (e), plant level LMA (f). F statistics and letters are given from Tukey tests, all traits were log-transformed.
